# Thiol-mediated Uptake of a Cysteine-containing Nanobody for Anti-Cancer Drug Delivery

**DOI:** 10.1101/2022.07.12.497993

**Authors:** Felix Goerdeler, Emelie E. Reuber, Jost Lühle, Sabrina Leichnitz, Anika Freitag, Ruslan Nedielkov, Heiko M. Möller, Peter H. Seeberger, Oren Moscovitz

## Abstract

The identification of tumor-specific biomarkers is one of the bottlenecks in the development of cancer therapies. Previous work revealed altered surface levels of reduced/oxidized cysteines in many cancers due to overexpression of redox-controlling proteins such as protein disulfide isomerases on the cell surface. Alterations in surface thiols can promote cell adhesion and metastasis, making thiols attractive targets for treatment. Only a few tools are available to study surface thiols on cancer cells and exploit them for theranostics. Here, we describe a nanobody (CB2) that recognizes B cell lymphoma in a thiol-dependent manner. CB2 binding strictly requires the presence of a non-conserved cysteine in the antigen-binding region and correlates with elevated surface levels of free thiols on B cell lymphoma compared to healthy lymphocytes. Nanobody CB2 can induce complement-dependent cytotoxicity against lymphoma cells when functionalized with synthetic rhamnose trimers. Lymphoma cells internalize CB2 in a thiol-mediated manner such that the nanobody can be used to deliver cytotoxic agents. Hence, surface thiols can be used as lymphoma biomarkers and targeted by thiol-binding nanobodies. Functionalization of internalizable CB2 is the basis for a range of diagnostic and therapeutic applications of this thiol-binding nanobody.

**TOC Graphic:** 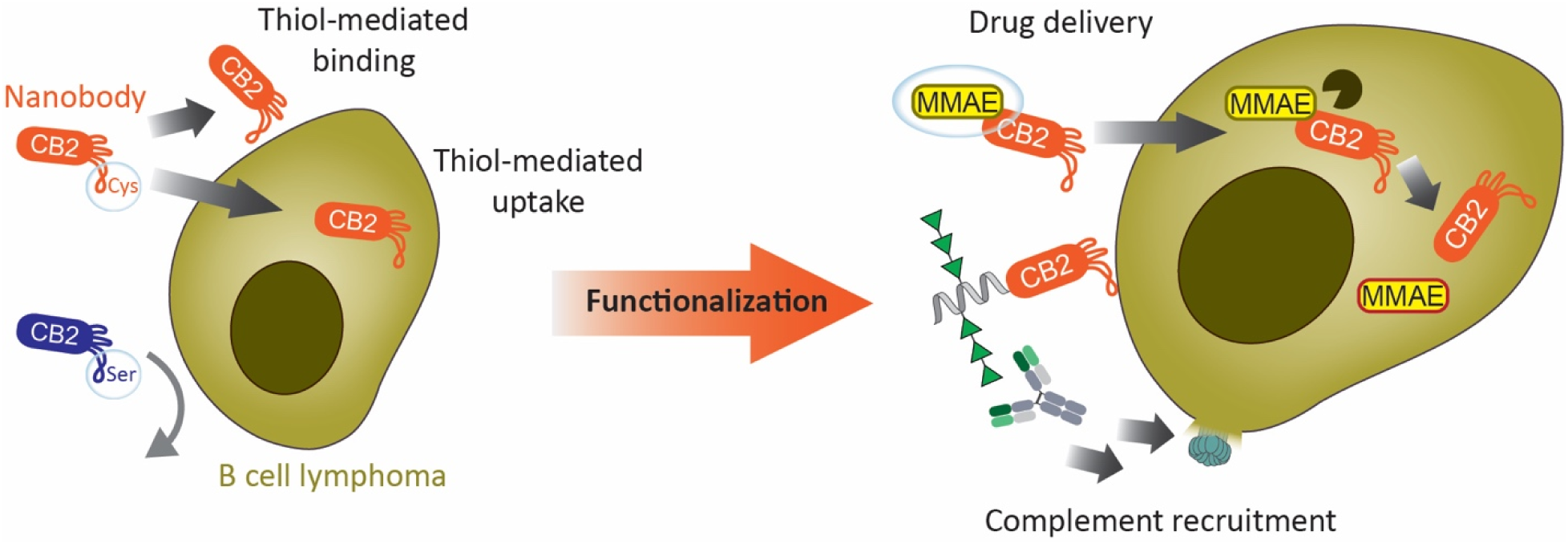

**Synopsis:** Nanobody CB2 specifically binds and internalizes into B cell lymphoma via thiol-based interactions. Functionalized CB2 can be used for complement recruitment or drug delivery to lymphoma cells.

## Introduction

Non-Hodgkin B cell lymphoma is one of the most common cancer types, that affects about 2% of men and women during their lifetime^1^. The main treatment strategy combines the antibody rituximab with chemotherapy, which in general affects also healthy cells^2^. Therefore, less harmful treatment options are highly desirable. The key challenge for developing safe and efficient cancer treatments is the identification of cancer-specific targets. In this respect, the extracellular cancer microenvironment has been an ample source for new biomarkers^3^. Previous work unveiled that the extracellular redox state of many cancers is significantly altered compared to healthy tissues^4–6^. Redox-controlling proteins, such as protein disulfide isomerases (PDIs) or the thioredoxin system, show high expression levels in various tumors^4,7,8^. Notably, the inhibition of PDIs reduces cancer proliferation, rendering PDIs promising therapeutic targets^7–9^. Thioredoxin and PDIs catalyze thiol-disulfide exchange reactions, leading to altered levels of reduced or oxidized protein disulfide bridges. For instance, recent evidence confirms that free thiol groups on the surface of breast cancer cells promote cell adhesion and metastasis^5^. The presence of reduced surface cysteines on cancer cells was also exploited as a vantage point for small thiol-functionalized compounds or peptides that bind to surface thiols resulting in cellular uptake^10–12^.

Nanobodies (Nbs) are the smallest antigen-binding fragments from heavy-chain-only antibodies, exclusively found in camelids (VHH) and cartilaginous fish (VNAR). Their single-domain nature allows straightforward expression in bacterial systems. Compared to conventional IgG antibodies (Abs) with approx. 150 kDa, their small size (approx. 15 kDa) enables Nbs to reach less accessible antigens while maintaining high affinity and stability. Functionalization conveys desired additional properties to Nbs, such as fluorescence, cytotoxicity, or cell internalization^13^. Introducing positive charges, either directly into the Nb framework or by fusing a charged peptide, leads to rapid cellular uptake of the engineered Nbs^14,15^.

In human blood, endogenous Abs against a variety of small molecules and glycans are highly abundant because these molecules are recognized as non-self by the immune system^16,17^. For example, rhamnose (Rha) is found in many bacterial polysaccharides, and humans produce anti-Rha Abs due to constant exposure to bacteria in their environment. To exploit these naturally occurring anti-Rha Abs for cancer therapy, cancer cells have been labeled with rhamnose using rhamnose-functionalized liposomes or antibodies^18,19^. Once cancer cells are rhamnose-tagged, anti-Rha Abs from human serum recognize the cells and activate downstream immune pathways, such as the complement cascade, leading to cancer cell death *in vitro* and *in vivo*^18,19^.

Here we describe CB2, a Nb discovered by serendipity that specifically binds to reduced surface cysteines upon which the complex internalizes. We show that CB2 binding and internalization correlate with higher surface levels of reduced cysteines on lymphoma cells compared to healthy lymphocytes and that CB2 can be easily functionalized for different applications.

## Results and Discussion

### CB2 recognizes several B cell lymphoma cell lines but not healthy lymphocytes

As part of a previous study for Nb generation^20^, we screened several Nb candidates for binding to antigens we had used to immunize an alpaca. In parallel, binding to several B cell lymphoma cell lines was tested by flow cytometry. By serendipity, we found one Nb candidate, named CB2, with surprising binding activity: While CB2 did not recognize any of the antigens used for immunization, it reliably bound to the B cell lymphoma cell lines SC-1, Raji, Jeko-1, DOHH-2, and SU-DHL-4, representing several subtypes of B cell lymphoma (Fig. 1). In contrast, healthy human peripheral blood mononuclear cells (PBMCs) were not recognized by CB2 (Fig. 1). To further characterize CB2 binding, we chose SC-1 (follicular lymphoma) as a model cell line. First, we determined the level of CB2 binding to SC-1 cells over time by flow cytometry and found no differences between incubation for 30 min, 1 h or 2 h (Fig. S1). On-cell affinity measurements revealed an apparent dissociation constant of CB2 in the low micromolar range (K_D_* = 6.3 ± 0.4 μM) (Fig. S1). Based on these findings, we were intrigued to determine the molecular basis of CB2’s specificity for lymphoma cells.

**Figure 1:**
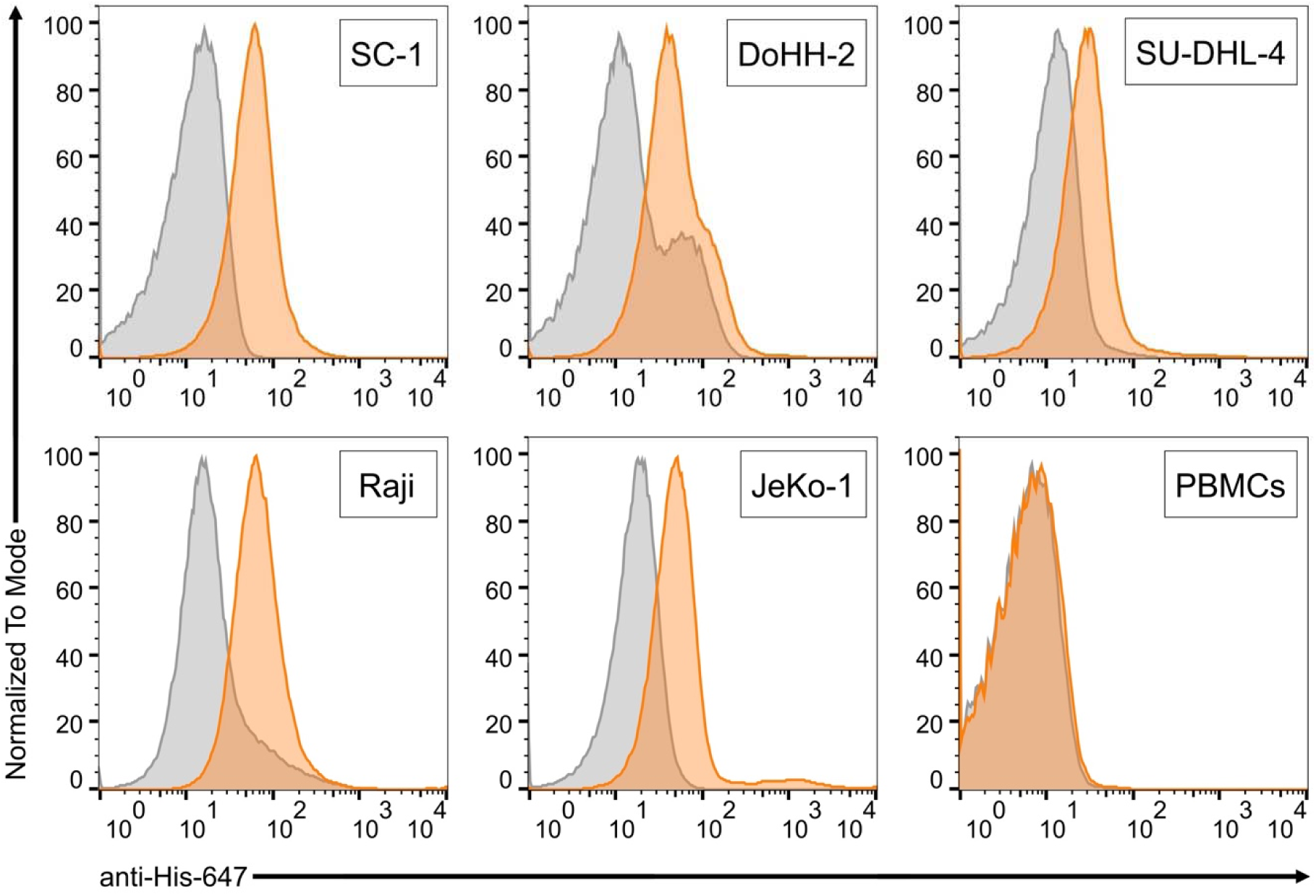
CB2 specifically binds several subtypes of B cell lymphoma. Flow cytometry binding assays with different lymphoma cell lines and healthy human PBMCs. Grey histograms show secondary antibody controls. Screened cells are SC-1 (Follicular lymphoma), DoHH-2 (Follicular lymphoma), SU-DHL-4 (Diffuse large B cell lymphoma), Raji (Burkitt’s lymphoma), JeKo-1 (Mantle cell lymphoma), and healthy human PBMCs.

### Lymphoma cells display higher levels of accessible surface thiol groups than healthy lymphocytes

During CB2 purification, a substantial part of the protein was obtained as dimers, that are not formed in the presence of reducing agents such as beta-mercaptoethanol or dithiothreitol (DTT) (Fig. S2A-B). We identified a cysteine residue in the complementarity-determining region 3 (CDR3) of CB2 at position 105 and speculated that CB2 interaction with cancer cells may be based on thiol-thiol interactions. Previous work revealed that the intra- and extracellular redox state of cancer cells is often altered in the course of cancer progression, for instance, due to PDI overexpression^4,5,7,8^. However, we could not find any quantitative data on the amounts of free thiol groups on lymphoma cells. Therefore, we set out to compare the level of free thiols on SC-1 cells and healthy lymphocytes using the thiol probe Alexa647-maleimide. After verifying by confocal microscopy that this probe is not internalized by cells and remains on the cell surface (Fig. S3), we quantified the surface thiol levels of cells by flow cytometry. Indeed, SC-1 lymphoma cells showed approx. five-fold increased fluorescence compared to healthy lymphocytes, suggesting higher levels of accessible surface thiol groups on lymphoma cells (Fig. 2A).

**Figure 2:**
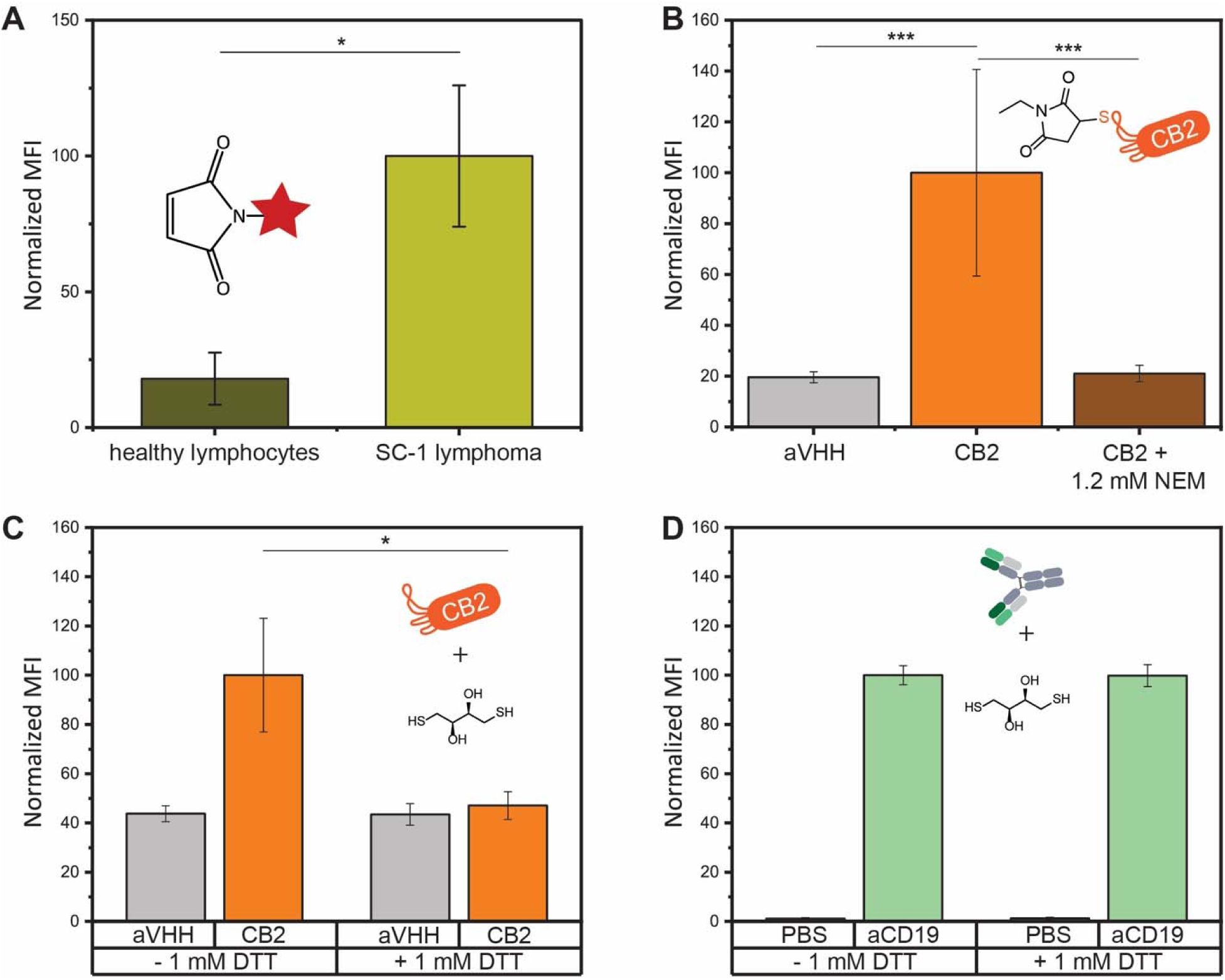
Altered surface thiol levels on lymphoma cells mediate CB2 binding. (A) Flow cytometry quantification of reduced surface thiols on lymphoma cells and healthy lymphocytes. (B) CB2 binding to SC-1 cells is completely lost after pre-incubation of CB2 with *N*-ethylmaleimide. (C-D) Binding to SC-1 cells in the presence or absence of 1 mM DTT. (C) CB2 binding is abolished in the presence of DTT. (D) The binding of the anti-CD19 Ab (positive control) is not affected by DTT. Values represent mean fluorescence intensity (MFI) from three independent experiments. Error bars represent the standard error of the mean (SEM). Differences were tested for significance using one-way ANOVA followed by Tukey’s *post-hoc* test with (*) p < 0.05, (**) p < 0.01, (***) p < 0.001.

### CB2 binding requires a reduced, accessible thiol group

To determine whether CB2 binding is mediated by thiol-thiol interactions, we performed a variety of inhibition experiments (Fig. 2B-D). When repeating CB2 binding assays in the presence of the reducing agent DTT, CB2 binding to SC-1 cells was completely abolished (Fig. 2C). The binding of anti-CD19 Ab, used as a positive control, remained unchanged, demonstrating that mildly reducing conditions do not interfere with Ab binding in general (Fig. 2D).

In addition, we also pre-incubated CB2 with the thiol quenching agent N-ethylmaleimide (NEM) before probing cell binding and found that NEM-treated CB2 lost binding to SC-1 cells, further highlighting the crucial role of a free thiol group for CB2 binding (Fig. 2B).

### Cysteine 105 is essential for CB2 binding

Encouraged by our result, we generated a C105S mutant of CB2 (hereon ^C105S^CB2) to examine the role of this specific residue in cell recognition (Fig. S2A). Indeed, ^C105S^CB2 failed to bind SC-1 cells when testing its activity in flow cytometry assays (Fig. 3A-B). To exclude the possibility that the mutation disrupts the structure of CB2, we acquired 1H-15N HSQC NMR spectra of 15N-labeled CB2 and ^C105S^CB2. The spectra showed only minimal shifts in the signals (Fig. S2C), indicating that the mutant retained the same folding as CB2. To further corroborate the flow cytometry data, we also incubated SC-1 cells with CB2 or ^C105S^CB2 and stained bound Nb with anti-6xHis-Alexa647 Ab for confocal microscopy (Fig. 3C-D). Indeed, CB2 binding was completely abolished for the C105S mutant, confirming our previous observation. Cysteine 105 hence plays an essential role in the recognition of SC-1 cells by CB2.

**Figure 3:**
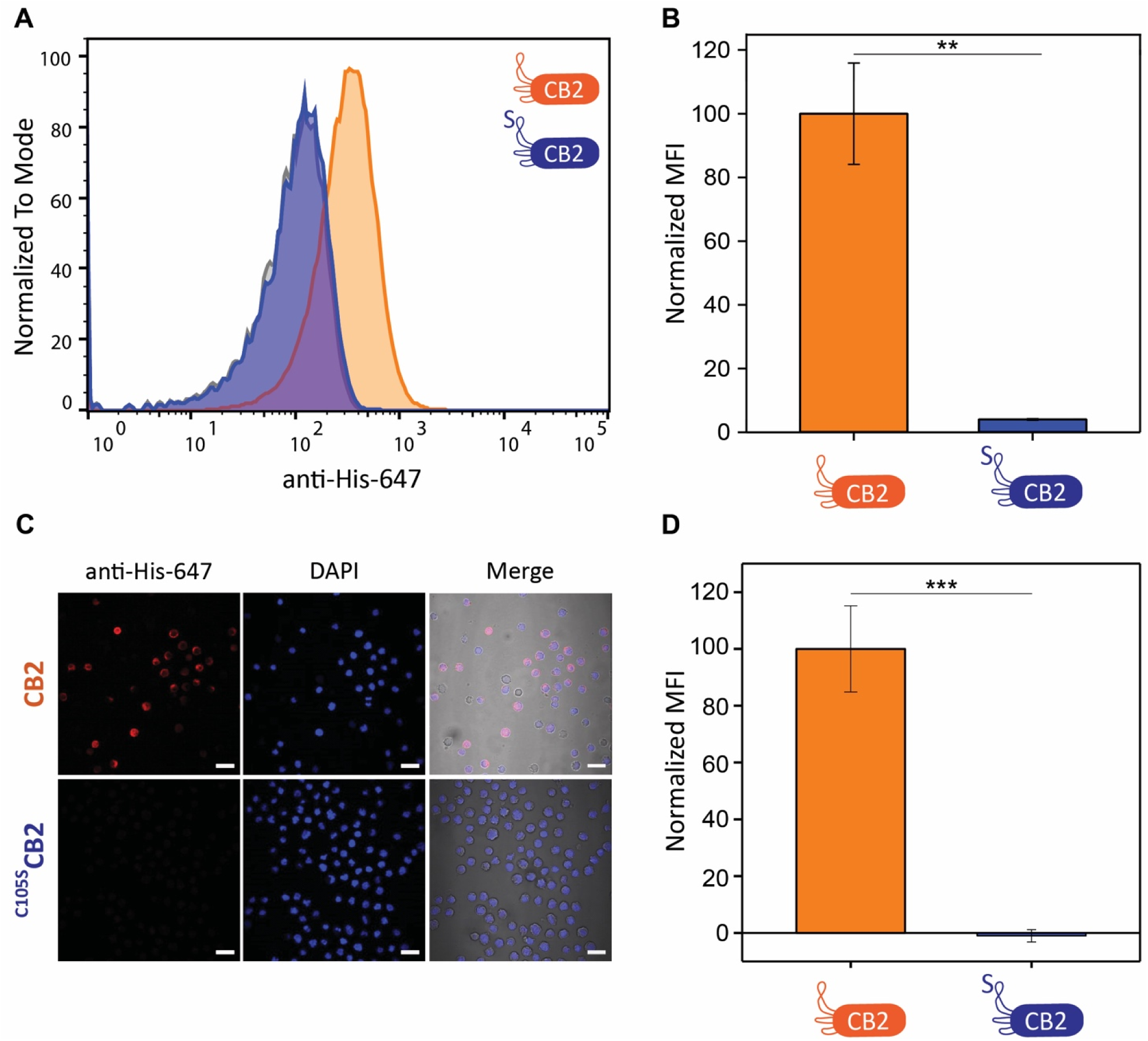
^C105S^CB2 is unable to bind B cell lymphoma. (A) Representative flow cytometry histograms of a binding assay to SC-1 cells. Secondary antibody control is displayed in grey. (B) Quantification of flow cytometry binding assays of CB2 and ^C105S^CB2 to SC-1 cells (N = 3). Normalized MFI is significantly reduced for ^C105S^CB2. (C) Confocal fluorescence microscopy images of SC-1 cells incubated with CB2 and ^C105S^CB2, respectively. Scale bars correspond to 20 μm. Red: anti-His-647, blue: DAPI, greyscale: transmission light. (D) Quantification of fluorescence microscopy images shown in A (N = 2 and n > 170 cells). Normalized MFI is significantly reduced in the case of ^C105S^CB2. Values represent mean ± SEM. Differences were tested for significance using one-way ANOVA followed by Tukey’s *post-hoc* test with (*) p < 0.05, (**) p < 0.01, (***) p < 0.001.

### Rhamnose-functionalized CB2 triggers complement activation against lymphoma cells

Due to the established role of rhamnose as an antibody-recruiting molecule (ARM), we reasoned that functionalization with rhamnose could enable CB2 to recruit Abs to lymphoma cells (Fig. 4A). In this model, Ab recruitment would trigger a downstream immune response by activating the complement cascade, ultimately leading to cancer cell death.

**Figure 4:**
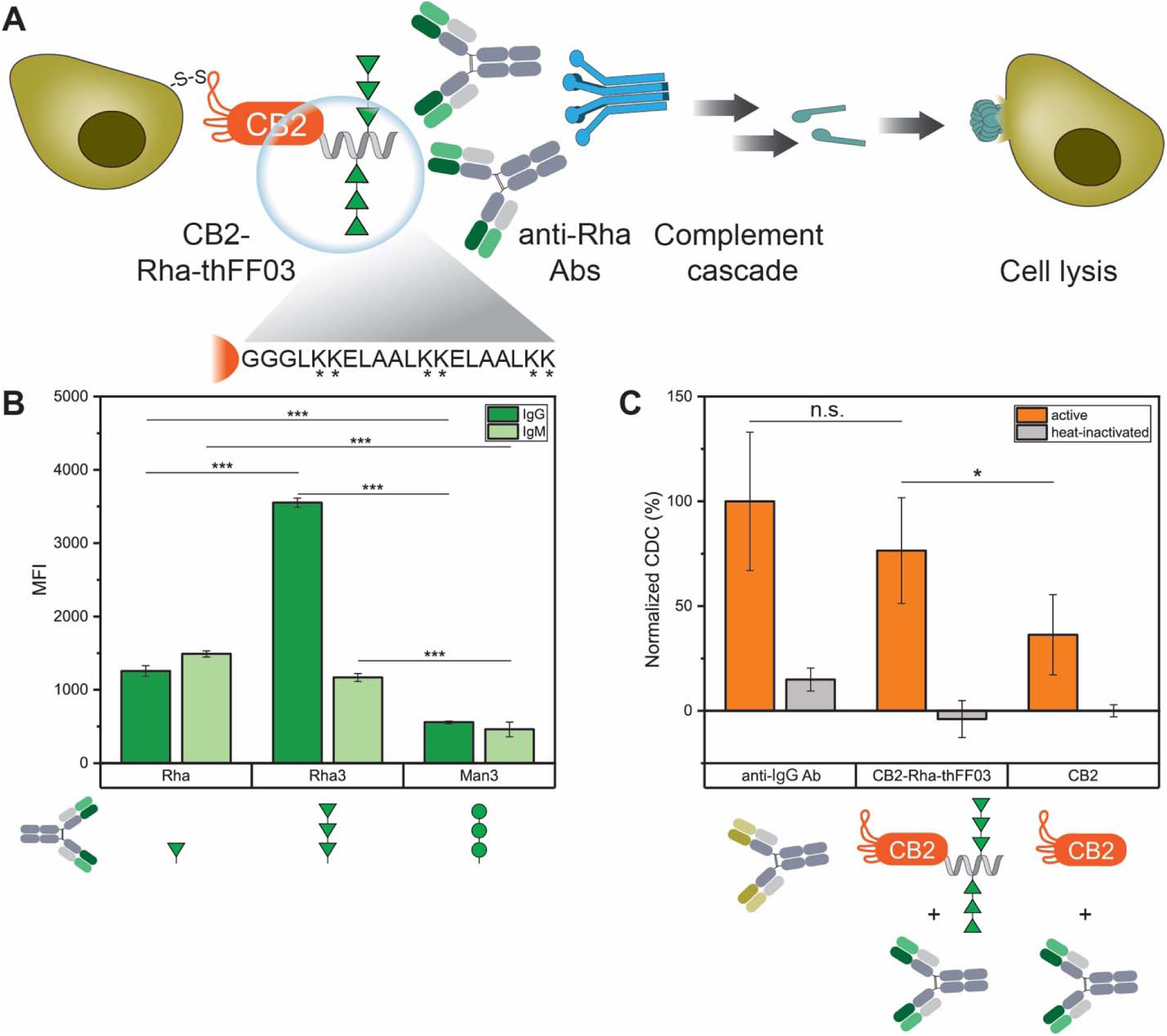
Rhamnose-functionalized CB2 induces CDC against B cell lymphoma. (A) Model for complement activation. CB2-Rha-thFF03 recruits endogenous anti-Rha Abs to lymphoma cells via its Rha moieties. Anti-Rha Abs activate the complement cascade, leading to cell lysis. Magnified: The thFF03 peptide contains six lysines for the chemical conjugation of Rha_3_. (B) Glycan array confirming the presence of anti-Rha Abs in human serum. Note that Rha_3_ is recognized by more IgGs isolated from human serum than Rha monomers. (C) Complement-dependent cytotoxicity assay. Note that CB2-Rha-thFF03 shows increased cytotoxicity compared to CB2. Anti-human IgG Ab was included as a positive control. Values are normalized to the positive control (100%) and represent mean ± SEM (N = 3), (*) p < 0.05, (**) p < 0.01, (***) p < 0.001.

When screening Abs isolated from healthy human serum on a synthetic glycan array for the presence of anti-Rha Abs, we could detect both IgG and IgM Abs binding to rhamnose (Fig. 4B). We then coupled glycylated rhamnose monosaccharides to the C-terminus of CB2 using sortase A (srtA) to test Ab recruitment to cells. However, we could not detect any complement-dependent cytotoxicity (CDC) of CB2-Rha on SC-1 cells (data not shown). Previous work on ARMs suggested that attaching a single ARM moiety might not be sufficient to recruit Abs, as anti-Rha Abs in human serum are formed due to natural exposure to Rha-decorated pathogens, and bacterial capsular polysaccharides often contain several Rha moieties in a row^19,21^. Since rha(α1-2)rha(α1-2)rha (hereon Rha_3_) forms part of the surface glycans of both *Klebsiella* and *Streptococcus*^22,23^, we compared serum IgG/IgM levels against a single Rha or Rha_3_ on glycan arrays. As a control, we measured the Ab titer against man(α1-2)man(α1-2)man (Man_3_). As expected, IgM/IgG titers both against Rha and Rha_3_ were significantly higher compared to Man_3_ (Fig. 4B). Interestingly, we could see a significant three-fold increase in fluorescence for IgGs against Rha_3_ compared to Rha, suggesting that more IgGs are present against Rha_3_ than against Rha (Fig. 4B).

Therefore, we designed the multivalent rhamnose conjugate CB2-Rha-thFF03 (Fig. S5), consisting of CB2 and a C-terminally coupled synthetic glycopeptide, decorated with several Rha_3_, and tested its effect on cells. Anti-human IgG Ab was used as a positive control for complement activation as it recognizes the abundant surface IgGs on B cell lymphoma. Indeed, CB2-Rha-thFF03 reliably induced complement activation against SC-1 cells, measured by an increase in dead SC-1 cells after incubation with CB2-Rha-thFF03, human anti-Rha Abs and active complement (Fig. 4C). Compared to the 100% cytotoxic effect of the positive control, CB2-Rha-thFF03 achieved 76% cytotoxicity. In contrast, unconjugated CB2 showed only 36% cytotoxicity, corresponding to unspecific complement activity (Fig. 4C). This demonstrates the ability of CB2-Rha-thFF03 to recruit Abs and complement factors to lymphoma cells.

Based on our observations, we recommend employing multivalent display of rhamnose multimers for antibody recruitment/complement activity assays. First, Rha_3_ showed much higher Ab response levels than Rha when investigating pooled human serum on a glycan array. Second, a single Rha_3_ attached to the C-terminus of CB2 was insufficient to recruit Abs to SC-1 cells, and we only observed complement-dependent cytotoxicity with the multivalent construct CB2-Rha-thFF03.

### CB2 binding causes cellular uptake via clathrin-mediated endocytosis

Since previous studies showed improved cellular uptake of thiol-functionalized peptides^24^, we hypothesized that CB2 could also be internalized by SC-1 cells. To test this hypothesis, we generated Cy3-functionalized CB2 and ^C105S^CB2 via a combination of srtA-mediated conjugation and click chemistry (Fig. S6A-B). After verifying that CB2-Cy3 conjugates still retain binding activity (Fig. S6D-E), we examined potential cellular uptake using confocal microscopy. Indeed, CB2-Cy3 is readily internalized by SC-1 cells as fluorescence accumulates inside the cytoplasm with CB2-Cy3 but not with ^C105S^CB2-Cy3, demonstrating the thiol-dependent nature of the binding mechanism which is a prerequisite for internalization (Fig. 5A-C, Fig. S6F-I). Interestingly, internalization is significantly reduced in the presence of clathrin-mediated endocytosis inhibitors dynasore or chlorpromazine, suggesting that CB2-Cy3 is endocytosed by SC-1 cells (Fig. 5A-C, Fig. S6F-I).

**Figure 5:**
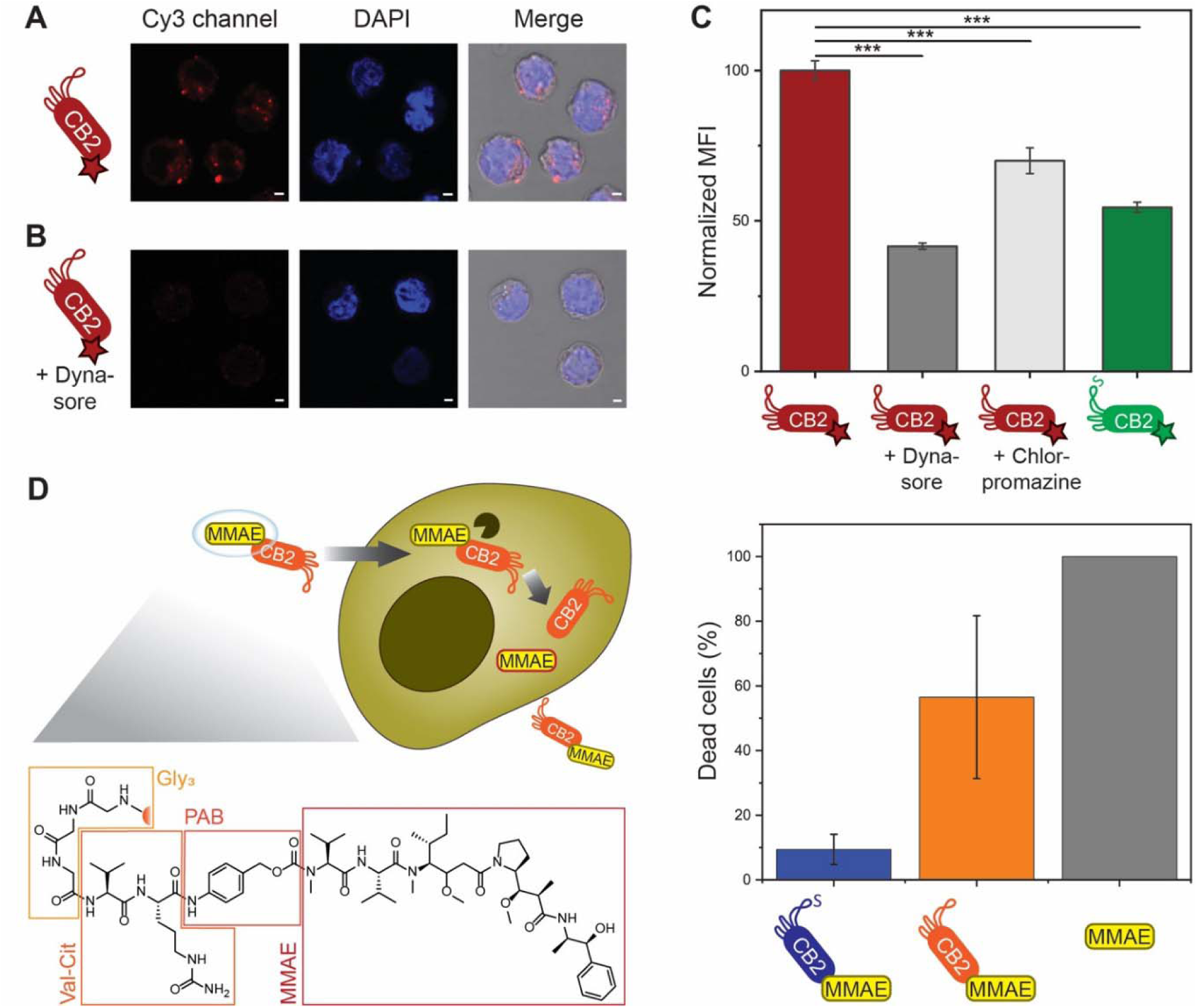
CB2 can be internalized by B cell lymphoma and used for drug delivery. (A-B) Exemplary images of CB2-Cy3 internalization by SC-1 cells in the absence (A) or presence (B) of the endocytosis inhibitor dynasore. Red: Cy3, blue: DAPI. Scale bar = 2 μm. (C) Quantified Cy3 fluorescence of SC-1 cells after 2 h of incubation with CB2-Cy3, CB2-Cy3 + dynasore, CB2-Cy3 + chlorpromazine, and ^C105S^CB2-Cy3. Note that both dynasore and chlorpromazine inhibit CB2 uptake and that no internalization is observed for the C105S mutant. Values represent mean ± SEM (N = 3 and n > 160 cells), (***) p < 0.001. (D) Drug delivery with internalizing CB2. Left panel: Proposed mechanism of action. MMAE is coupled to CB2 via a cleavable linker using srtA. Val-Cit = valine-citrulline, PAB = para-aminobenzylcarbamate, MMAE = monomethyl auristatin E. MMAE is released inside the cell by cathepsin (black) cleaving between val-cit. Right panel: Cytotoxicity assay with SC-1 cells. Dead cells (%) after 48 h with 10 nM of CB2-MMAE, ^C105S^CB2-MMAE, or MMAE. Note that only CB2-MMAE and MMAE alone show considerable cytotoxic activity. Cytotoxicity was measured using the CCK8 kit and normalized to 100%, and 0% based on positive (TritonX) and negative (unconjugated CB2) controls, respectively. Values represent mean ± SEM (N = 3).

### CB2 internalization can be exploited for drug delivery

To determine whether CB2 internalization can be used for potential therapeutic application, the srtA reaction was used to couple the antineoplastic agent monomethyl auristatin E (MMAE) to the C-terminus of CB2 (Fig. S7A-B). Due to its high potency and resulting off-target toxicity, MMAE cannot be administered alone but only when conjugated to an antibody or nanobody conveying target specificity^25^. The commercially available MMAE construct for sortase-mediated conjugation consists of the active compound and a valine-citrulline linker that is cleaved by intracellular cathepsines, thereby releasing the drug inside the cell (Fig. 5D).

After confirming that CB2-MMAE retains binding activity (Fig. S7C), we incubated lymphoma cells with 10 nM of MMAE, CB2-MMAE, ^C105S^CB2-MMAE or non-conjugated CB2 and measured cell viability after 48 h (Fig. 5D). Cells treated with 1% Triton X-100 or unconjugated CB2 served as positive (100% dead cells) and negative (0% dead cells) control, respectively. After 48 h, CB2-MMAE had killed ∼60% of cells compared to 100% dead cells observed with unconjugated MMAE. In contrast, ^C105S^CB2-MMAE killed only ∼10% of cells. (Fig. 5D). These results demonstrate that CB2 can indeed be used to deliver cytotoxic agents inside lymphoma cells, rendering CB2-MMAE conjugates a promising tool to circumvent MMAE’s high off-target toxicitiy.

Interestingly, CB2 internalization is observed even though we could not detect changes in the level of surface-bound CB2 within the time scale of CB2 internalization (Fig. S1). Compared to surface-bound CB2, the amount of internalized CB2 might be too less to be detected by our approach. However, the fact that CB2 binding and internalization are observed within a similar time scale suggests that CB2 targets distinct populations of membrane proteins. In this model, some protein targets remain on the surface while others get endocytosed upon CB2 binding, allowing for extra- and intracellular targeting of lymphoma cells with CB2. Indeed, Rha_3_-functionalized CB2 reliably recruited anti-Rha Abs to the surface of SC-1 cells, thereby inducing complement-dependent cytotoxicity. At the same time, MMAE-functionalized CB2 was able to deliver MMAE to the cytosol of SC-1 cells, causing drug-induced cytotoxicity.

## Conclusions

With CB2, we describe the first nanobody that recognizes and internalizes into lymphoma cells in a thiol-dependent manner. We demonstrate that CB2’s interaction with B cell lymphoma requires the presence of a free thiol group at cysteine 105. Specificity arises from increased surface levels of reduced thiol groups on B cell lymphoma compared to healthy lymphocytes. Finally, the thiol-dependent nature of CB2 binding also leads to CB2 uptake by B cell lymphoma. We show that CB2 internalization opens up a vast array of potential applications for cancer diagnostics and therapeutics. In addition to drug delivery, a diagnostic use of CB2 seems also promising as fluorescently labeled CB2 could be directly used for tumor imaging *in vivo*. Bispecific constructs are conceivable, combining CB2 with a nb against other lymphoma biomarkers such as CD19. This could potentially decrease off-target binding and, at the same time, lead to tumor inhibition via receptor internalization. Finally, CB2 could be employed to develop chimeric antigen receptor (CAR) T cells, rendering it applicable for immunotherapy of B cell lymphoma.

## Acknowledgments

Generous financial support by the Max Planck Society is gratefully acknowledged. This work was supported by Deutsche Forschungsgemeinschaft (RTG2046 for F.G. and J.L., CRC1449 for S.L.). The authors thank Prof. Helge Ewers for providing the sortase expression plasmid. The authors thank Dr. Peter Sondermann for reviewing the manuscript and providing helpful feedback.

