## Supplemental information for "Thiol-mediated Uptake of a Cysteine-containing Nanobody for Anti-Cancer Drug Delivery"

#### Experimental Section

1. Nanobody expression and purification
2. Site-directed mutagenesis of CB2
3. NMR measurements of CB2 and $^{C105S}$CB2
4. Cell culture
5. Nanobody binding assays and determination of apparent affinity
6. Thiol-binding inhibition assays
7. Determination of surface thiol levels
8. Synthesis of $Rha_3$
9. $Rha_3$ conjugation to CB2
10. Glycan array
11. Complement activation assay
12. Cy3 conjugation to CB2
13. Internalization assay
14. MMAE conjugation to CB2
15. Drug delivery assay

#### List of Figures

1. Apparent affinity and time course of CB2 binding to SC-1 cells
2. CB2 dimerization/mutant and NMR spectra
3. SC-1 cell staining with Alexa647-maleimide
4. NMR spectra of $Rha_3$
5. $Rha_3$ conjugation to CB2
6. Cy3 conjugation to CB2 and internalization controls
7. MMAE conjugation to CB2

### Experimental Section

No unexpected or unusually high safety hazards were encountered.

#### Nanobody expression and purification

The sequence of CB2 was synthesized by Synbio Technologies (New Jersey, USA) with a C-terminal glycine-serine linker ($[G_4S]_3$) followed by a sortag (LPETG) and a $His_6$ tag in a pET-28b(+) vector. The plasmid was transformed into *E. coli* SHuffle cells (NEB, Ipswich, MA, USA) and the protein was expressed in TB medium via induction with 0.5 mM IPTG at 16°C overnight. Bacteria were lysed using a French press and CB2 was purified using gravity nickel-NTA chromatography followed by size exclusion chromatography on an ÄKTApurifier with a Superdex 75 16/600 column (GE Healthcare). All steps of nanobody purification were validated by SDS-PAGE. CB2 was purified in the presence of 0.75 mM dithiothreitol to prevent dimerization.

#### Site-directed mutagenesis of CB2

The $^{C105S}$CB2 mutant was generated in-lab via overlap extension PCR as described elsewhere[1]. Briefly, forward and reverse primers containing the mutation were ordered from IDT (Coralville, IA, USA) and two separate PCRs were performed using each primer and the corresponding T7 standard primer. In a second PCR, both fragments were joined via overlap extension and the final product was cloned into the pET-28b(+) backbone. Expression of the mutant was carried out as described before.

#### NMR measurements of CB2 and $^{C105S}$CB2

For NMR measurements, CB2 and $^{C105S}$CB2 were expressed in labeled M9 minimal medium containing 1 g/L $^{15}N$-$NH_4Cl$ (Merck), 1 g/L NaCl, 3 g/L $KH_2PO_4$, 6 g/L $Na_2HPO_4$, pH 7.4, 2 mM $MgSO_4$, 100 µM $CaCl_2$, vitamin mix and 100 mg/L $^{15}N$-Celtone (Cambridge Isotope Laboratories) via induction with 0.5 mM IPTG at 16°C, overnight. The bacteria were lysed, and proteins purified as described above. After purification, the proteins were concentrated using Amicon-4 Ultra (MWCO = 3 kDa) and transferred to PBS buffer supplemented with 0.75 mM DTT, 8 mM $NaN_3$, 0.5 mM DSS and 5% $D_2O$. The final concentration in the NMR samples was 160 µM and 70 µM for CB2 and $^{C105S}$CB2, respectively.

NMR measurements were performed on a Bruker NEO 500 MHz NMR spectrometer equipped with a TCI cryogenic probe head with Z gradient. $^{1}H$-$^{15}N$ HSQC spectra were acquired using a standard Bruker pulse sequence with an acquisition time of 140 ms and 256 increments, with 16 scans for CB2 and 160 scans for $^{C105S}$CB2. Total measurement time was 1 h 20 min and 13 h 30 min for CB2 and $^{C105S}$CB2, respectively. All NMR experiments were performed at 298 K. The spectra were processed and analyzed using Bruker Topspin 4.1.4.

#### Cell culture

All B cell lymphoma cell lines, except for SC-1, were a kind gift by Uta Hoepken and Armin Rehm (MDC Berlin). SC-1 cells were acquired from DSMZ. All B cell lymphoma cell lines were cultured in RPMI + 10% fetal calf serum + 2 mM glutamine + 1 mM sodium pyruvate + 1x penicillin/streptomycin at 37 °C, 5% $CO_2$. Cells were diluted into fresh medium every 2-3 days. All cell lines were tested for mycoplasma contamination on a monthly basis.

#### Nanobody binding assays and determination of apparent affinity

For flow cytometry screenings, one million cells per sample of the respective cell line or healthy human peripheral blood mononuclear cells (PBMCs) were stained with 24 µM nanobody solution or PBS only (negative control). Secondary staining was conducted with 50 µl anti-6X His Tag ATTO 647N conjugated antibody (Rockland Immunochemicals, Inc.) in a 1:750 dilution. Stained samples were measured on a FACSCanto™ II device (BD Biosciences). Events were gated for single cells and the median fluorescence intensity (MFI) of the APC-A signal was exported using the software FlowJo (V10.8.1). MFIs were subtracted with the negative control for background correction. MFIs from three independent experiments were determined and compared to examine differences in cell binding. For the determination of apparent affinity via flow cytometry, SC-1 cells were incubated as described above with different concentrations of CB2 (0.05-22 µM). Bound fractions of cells were determined with the software FlowJo and for each concentration, the relative binding was calculated based on the maximum binding at saturation. Data from three independent experiments was plotted and fitted using a 2-parameter sigmoidal regression model with the following equation:

$$y = \frac{1}{1 + \exp\left(\frac{-x + x_0}{b}\right)}$$

Apparent affinity ($K_D$*) was exported from the fit by determination of $x_0$. Error bars of the data points as well as the error (±) of the $K_D$* values correspond to standard errors.

For binding studies using confocal microscopy, two million cells per sample were washed once with PBS and resuspended in 50 µL of CB2 or $^{C105S}$CB2 (24 µM) in PBS. The negative control was resuspended in PBS. Cells were incubated 1 h at RT and shaking at 400 rpm. After three washes with PBS, cells were incubated with 50 µL anti-6X His Tag Atto 647N (1.3 µg/mL, Rockland Inc.) for 1 h at RT in the dark at 400 rpm. Cells were washed three times with PBS and resuspended in 1 mL PBS. They were settled on 25 mm coverslips in 6-well plates for 30 min in the dark at RT. Afterwards the supernatants were aspirated carefully and 750 µL of 4% paraformaldehyde (PFA) + 0.2% glutaraldehyde (GA) were added to each well for fixation. Samples were fixed at RT for 15 min in the dark. Afterwards, the coverslips were washed three times with 1 mL PBS, flipped onto a microscopy glass slide prepared with 50 µL of Roti®Mount FluorCare DAPI mounting solution (Carl Roth, Darmstadt, Berlin) and sealed with transparent nail polish. The slides were examined and imaged under a LSM700 laser scanning confocal microscope (Zeiss). The images were analyzed using Fiji (Fiji Is Just ImageJ) by manually selecting cells and then measuring their fluorescence intensity (mean grey value). The mean fluorescence intensity (MFI) was normalized using the following equation:

$$Normalized\ MFI\ (x) = \frac{MFI(x) - MFI\ (PBS)}{MFI(CB2) - MFI\ (PBS)}$$

The statistical analysis was performed with the software GraphPad Prism 9.3.1 (GraphPad Software, Inc.).

##### Thiol-binding inhibition assays

For DTT inhibition experiments, incubation of CB2 and SC-1 cells was carried out in the presence of 1 mM DTT. Anti-CD19-PE (1:100, BioLegend) was used as positive control Ab, as DTT should not affect binding of this Ab to B cells. For NEM inhibition experiments, CB2 was pre-incubated with 1.2 mM NEM for 2 h at RT. Excess NEM was washed out thrice using an Amicon Ultra Centrifugal Filter (cut-off = 10 kDa) and NEM-pre-treated CB2 was used for incubation with SC-1 cells. Nanobody binding was detected with MonoRab™ Rabbit Anti-Camelid VHH Cocktail iFluor 647 (1:500, GenScript).

#### Determination of surface thiol levels

Peripheral blood mononuclear cells (PBMCs) were freshly isolated as described elsewhere[2]. One million PBMCs or SC-1 cells were pelleted and incubated with AF647-maleimide (1:1,000, Jena Bioscience) or PBS for 15 min on ice in the dark[3]. Cells were washed thrice with PBS and analyzed by flow cytometry. An additional sample was prepared the same way and confocal microscopy was used to confirm that AF647-maleimide only stains the cell membrane but not intracellular compartments. The mean fluorescence intensity from three independent FACS experiments was quantified and background-corrected. Differences between healthy lymphocytes and SC-1 cells were tested for significance using student's t-test in Origin.

#### Synthesis of $Rha_3$

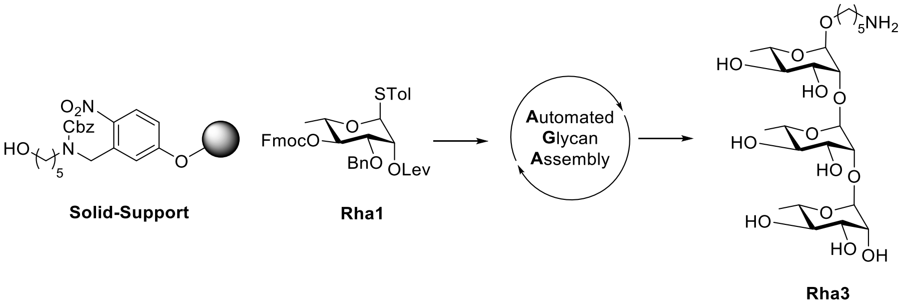


| Repeat | Building Blocks | Modules | Notes | |
|---|---|---|---|---|
| 3x | | **I** – Acidic Wash | | |
| | Rha1 (2 x 5.0 equiv.) | **IIa** – Glycosylation with thioglycoside – 2 cycles | -20 °C ($T_1$)<br>0 °C ($T_2$) | 10 min ($t_1$)<br>30 min ($t_2$) |
| | | **III** – Capping<br>**IVc** – Lev Deprotection | | |

The automated synthesis of $Rha_3$ was performed on a home-built synthesizer developed at the Max Planck Institute of Colloids and Interfaces using modules reported earlier[4]. Protected $Rha_3$ (35 mg, 0.022 mmol, crude yield: 80%) was obtained as a colorless oil after photo-cleavage from solid support with a UV-150 Medium-Pressure Mercury Lamp (arc length 27.9 cm, 450 W) surrounded by a long-pass UV filter (Pyrex, 50% transmittance at 305 nm). Deprotection of $Rha_3$ was achieved by methanolysis and hydrogenolysis. Briefly, to a solution of protected oligosaccharide in MeOH:$CH_2Cl_2$ (2 mL, 1:1), sodium methoxide (0.5 M solution in MeOH, 2.2 equiv. per ester group) was added. The mixture was stirred at room temperature for 2 h. Then Amberlite IR-120 ($H^+$ form) was added to quench. After neutralization, the reaction mixture was filtered and the solvent was removed in vacuo. The crude compound was dissolved in 4 mL of EtOAc:t-BuOH:$H_2O$ (2:1:1). Pd/C (10%) was added to the solution and the suspension was stirred in a $H_2$ bomb with 60 psi pressure over night. The insoluble material was removed by a CHROMAFIL ®Xtra, RC 0.45 syringe filter. The solid was washed once with t-BuOH and several times with water. The filtrate was collected and concentrated in vacuo. Crude products were dissolved in water and analyzed/purified using analytical/preparative HPLC. A Thermo-Scientific Hypercarb column (150 mm x 4.60 mm I.D.) was used for analytical RP-HPLC with a flow rate of 0.70 mL/min with water (0.1% $HCO_2H$)/acetonitrile as eluents (100% $H_2O$ (0.1% $HCO_2H$) for 5 min, 0 → 30% acetonitrile in $H_2O$ (0.1% $HCO_2H$) over 30 min, 30 → 100% acetonitrile in $H_2O$ (0.1% $HCO_2H$) over 5 min, 100% acetonitrile for 5 min). This yielded deprotected, pure $Rha_3$ (6 mg, 0.011 mmol, 41%) as a white solid after lyophilization. NMR spectra confirming the purity of the compound can be found in Fig. S4.

**$^1$H NMR** (700 MHz, $D_2O$): δ 5.11 (s, 1H), 4.99 (s, 1H), 4.87 (s, 1H), 4.09 (ddd, J = 11.5, 3.4, 1.7 Hz, 2H), 3.95 – 3.83 (m, 3H), 3.82 – 3.67 (m, 5H), 3.56 (dt, J = 10.0, 6.1 Hz, 1H), 3.53 – 3.43 (m, 3H), 3.01 (t, J = 7.6 Hz, 2H), 1.76 – 1.61 (m, 4H), 1.52 – 1.41 (m, 2H), 1.31 (d, J = 6.3 Hz, 3H), 1.30 (d, J = 6.2 Hz, 3H), 1.28 (d, J = 6.2 Hz, 3H) ppm. **$^{13}$C NMR** (176 MHz, $D_2O$): δ 102.2, 100.9, 98.4, 78.5, 78.3, 72.1, 72.1, 72.0, 70.2, 70.1, 70.0, 69.8, 69.3, 69.2, 68.8, 67.7, 39.4, 28.0, 26.6, 22.5, 16.7, 16.6, 16.6 ppm. **HRMS** (QToF): Calcd for $C_{23}H_{44}NO_{13}$ $[M + H]^+$ 542.2807; found 542.2813.

#### $Rha_3$ conjugation to CB2

$Rha_3$ was coupled to CB2 in a two-step reaction to achieve multivalent display while maintaining CB2 binding activity. First, $Rha_3$ was coupled to the synthetic peptide thFF03 (sequence: GGGLKKELAALKKELAALKK, synthesized by ProteoGenix). The peptide thFF03 is a truncated, glycylated version of the previously published hFF03[5] and was chosen because its many lysines allow chemical conjugation, while the N-terminal glycines make it amenable to srtA conjugation. $Rha_3$ with an aminopentanol linker was conjugated to thFF03 as described elsewhere[6]. Briefly, 1 eq. of $Rha_3$ was mixed with 10 eq. of homobifunctional adipic acid *p*-nitrophenyl diester in 300 µL DMSO + 25 µL pyridine + 10 µL triethylamine and stirred for 3 h at 300 rpm, RT. After lyophilization, esterified $Rha_3$ was washed 3x with a 1:1 mixture of diethyl ether and dichloromethane and 3x with a 1:4 mixture of the same solvents until uncoupled linker was no longer detected in the wash fractions by UV light. The absence of glycan in the wash fractions was confirmed by thin-layer chromatography. Then, 6 eq. of esterified $Rha_3$ was mixed with 1 eq. of thFF03 in 140 µL conjugation buffer (0.1 M sodium phosphate pH 8) and stirred for 24 h at 70 rpm, RT. Successful conjugation and degree of loading was assessed by MALDI and the glycopeptide (hereon Rha-thFF03) was purified by HPLC on a C18 column.

In a second step, Rha-thFF03 was conjugated to CB2 via srtA coupling. 15 nmol of CB2, 3.75 nmol of srtA, 406 nmol of Rha-thFF03 and 40 µL of equilibrated HisPur Ni-NTA resin (ThermoFisher) were mixed in 300 µL of srtA buffer (150 mM NaCl, 10 mM CaCl2, pH 7.5) and incubated rotating for 30 min at 4 °C. Resin was pelleted (700 g, 2 min) and the supernatant was checked for the presence of CB2-Rha-thFF03 by SDS-PAGE and MALDI and brought into PBS via Amicon® Ultra centrifugal filter units (MWCO = 10 kDa).

##### Glycan array

Human IgG/IgM Abs were purified from human serum (Seraclot, PanBiotech) via protein A/G column (ThermoFisher). The glycans were dissolved at 0.1 mM in 50 mM sodium phosphate buffer pH 8.5 and printed in 64 identical fields to NHS activated hydrogel glass slides (CodeLink slides, Surmodics) using a non-contact S3 microarray spotter (Scienion, Berlin, Germany). After incubation overnight in a humidified box, the remaining NHS groups of the slides were quenched with ethanolamine. The slides were blocked with 1% (w/v) bovine serum albumin (BSA) in phosphate buffered saline (PBS) and a 64 well incubation gasket (FlexWell Grid, Grace Bio Labs) was attached. The slides were incubated with human serum (Sigma-Aldrich, Darmstadt, Germany) or purified Abs diluted 1:100 in 1% BSA-PBS for 1 h at 37°. After three washes with PBS containing 0.1% (v/v) Tween-20 (PBS-T) the slides were incubated with goat anti-human IgG (H+L) AlexaFluor 647 (Invitrogen, Cat. A21445) and goat anti-human IgM (µ chain) AlexaFluor

488 (Invitrogen, Cat. A21215) diluted 1:400 for 1 h at 37°C. The slides were washed twice with PBS-T. After removing the gasket, the slides were washed once with PBS and once with water. The dried slides were scanned with a GenePix 4300A microarray scanner (Molecular Devices). Intensities were evaluated with GenePix Pro 7 (Molecular Devices). The statistical analysis was performed with the software GraphPad Prism 9.3.1 (GraphPad Software, Inc.).

##### Complement activation assay

Human IgG/IgM Abs (0.2 mg/mL) were mixed with CB2 or CB2-Rha-thFF03 (0.4 mg/mL) and incubated for 1 h at RT. Two aliquots of one million SC-1 cells each were incubated with the CB2 or CB2-Rha-thFF03 Ab mixtures for 1 h at RT, 400 rpm. Two samples incubated with goat anti-human IgG Ab (0.24 mg/mL, Invitrogen) were included as positive controls because this Ab binds to the surface IgGs of B cell lymphoma. After 1 h incubation, cells were pelleted, resuspended in 50 µL of active or heat-inactivated (30 min at 56 °C) rabbit complement (Cedarlane) and incubated for 2 h at 37 °C. Cells were again pelleted, resuspended in 200 µL of 7-AAD (1:100, BioLegend) and cell viability was measured on a FACSCanto II (BD Bioscience).

##### Cy3 conjugation to CB2

Cy3 conjugates were prepared as described previously[7]. 100 µM CB2 or $^{C105S}$CB2, 25 µM srtA and 3 mM dibenzocyclooctyne-NH2 (DBCO-amine) were mixed in 1.2 mL srtA buffer (150 mM NaCl, 10 mM CaCl2, pH 7.5) for 30 min at 4° C with 180 µL Ni-NTA resin on a rotation wheel for srtA-mediated functionalization of sortagged nanobodies with dibenzocyclooctyne-NH2 (DBCO-amine). The supernatant containing CB2-DBCO or $^{C105S}$CB2-DBCO was collected, and remaining DBCO-amine was removed using an PD MidiTrap™ G-10 column (Cytiva, Freiburg, Germany). Nanobody-DBCO conjugate was concentrated in Amicon® Ultra centrifugal filter units (MWCO = 3 kDa) at 4° C, 3,200 g and concentrations were determined by NanoDrop® ND-1000. The successful conjugation and purity were analyzed on an SDS-Page and the conjugate was stored at 4° C until proceeding with the click-reaction.

40 µM CB2-DBCO or $^{C105S}$CB2-DBCO conjugate was mixed with 80 µM Cy3-azide (Jena Bioscience) in click reaction buffer (20 mM HEPES pH 7.5, 300 mM NaCl and 10%

glycerol) and incubated for 17 h. Unbound reaction partners were removed by using an PD MidiTrap™ G-10 column (Cytiva, Freiburg, Germany). Successful labeling was verified by reducing SDS-PAGE and FACS measurement. The conjugate (CB2-Cy3 or $^{C105S}$CB2-Cy3) was stored at 4°C and used for internalization assays within a week.

#### Internalization assay

1.5 million SC-1 cells were incubated with 100 µL of CB2-Cy3 or $^{C105S}$CB2-Cy3 (0.5 mg/mL) for 2 h at RT, 400 rpm in the dark. Cells were pelleted, washed once with 1 mL ice-cold acidic wash buffer (100 mM glycine, 150 mM NaCl, pH 2.2), to remove surface-bound protein, and twice with PBS. Cell pellets were resuspended in 250 µL each and settled onto 12 mm coverslips for 30 min at RT in the dark. The suspension was carefully aspirated, and cells were fixed with 250 µL fixative (4% paraformaldehyde + 0.2% glutaraldehyde in PBS) for 15 min at RT in the dark. Coverslips were washed thrice with PBS, mounted onto a drop of Fluoromount G (ThermoFisher), sealed and stored in the dark till imaging.

For endocytosis inhibition assays, cells were pre-treated for 30 min with 80 µM dynasore or chlorpromazine and then incubated with CB2-Cy3 in the presence of the respective inhibitor.

#### MMAE conjugation to CB2

MMAE was conjugated to CB2 via srtA coupling. 30 nmol of CB2, 7.5 nmol of srtA, 900 nmol of $Gly_3$-Val-Cit-PAB-MMAE (MedChemExpress, in DMSO) and 80 µL of equilibrated HisPur Ni-NTA resin (ThermoFisher) were mixed in 600 µL of srtA buffer (150 mM NaCl, 10 mM CaCl2, pH 7.5) and incubated rotating for 30 min at 4 °C. Resin was pelleted (700 g, 2 min) and the supernatant was checked for the presence of CB2-MMAE by SDS-PAGE and MALDI. The supernatant was dialysed with 3x 2 L PBS to remove any uncoupled drug and concentrated via Amicon® Ultra centrifugal filter units (MWCO = 10 kDa) before estimating the protein concentration via microBCA kit (ThermoFisher). Drug conjugates were filtered through 0.22 µm and stored at 4°C.

#### Drug delivery assay

In a 96-well plate, triplicates of CB2-MMAE, $^{C105S}$CB2-MMAE and MMAE were prepared with 10 µL at 190 nM each. Wells with unconjugated CB2 and 0.1% Triton-X100 were included as negative and positive controls, respectively. SC-1 cells were resuspended in complete medium without phenol red, and 30,000 cells in 180 µL medium were added to each well. As another control, three empty wells were filled with medium without cells. The plate was incubated for 48 h at 37°C, 5% $CO_2$. Next, 10 µL of WST-8 reagent (from CCK8 kit, Abcam) were added to each well and, after another incubation for 3 h at 37°C, 5% $CO_2$, the absorbance of the plate was measured at 460 nm with a Tecan Infinite plate reader.

#### Supplementary Figures

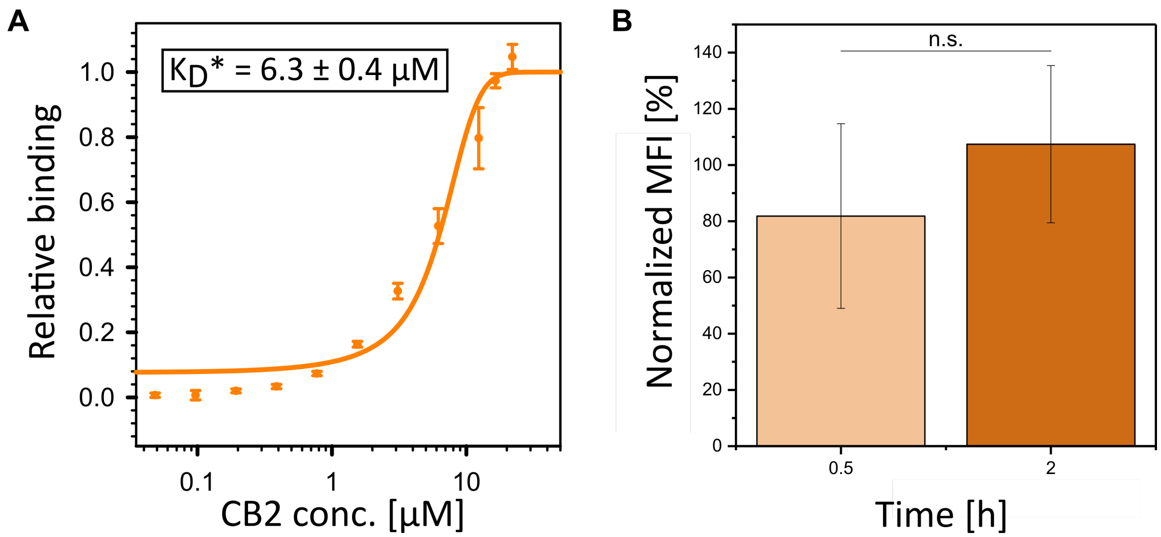


**Figure S1: (A) Apparent affinity of CB2 to SC-1 cells**. On-cell affinity was determined via flow cytometry incubating the cells with different concentrations of CB2. Data from three independent experiments is shown. Error bars of the data points and error (±) of the apparent dissociation constant ($K_D$*) correspond to standard errors. **(B) Time course of CB2 binding to SC-1 cells**. Binding was tested via flow cytometry after different time points of incubation with CB2 (0.5 h, 1 h, 2 h) to show that CB2 is still partially accessible on the cell surface after prolonged incubation. MFIs were normalized to the MFI of the 1 h time point (standard assay condition). There was no significant change in binding, indicating a similar surface retention of CB2 after prolonged incubation. Data from three individual experiments is shown. Error bars represent standard errors. Significance was tested via t-test.

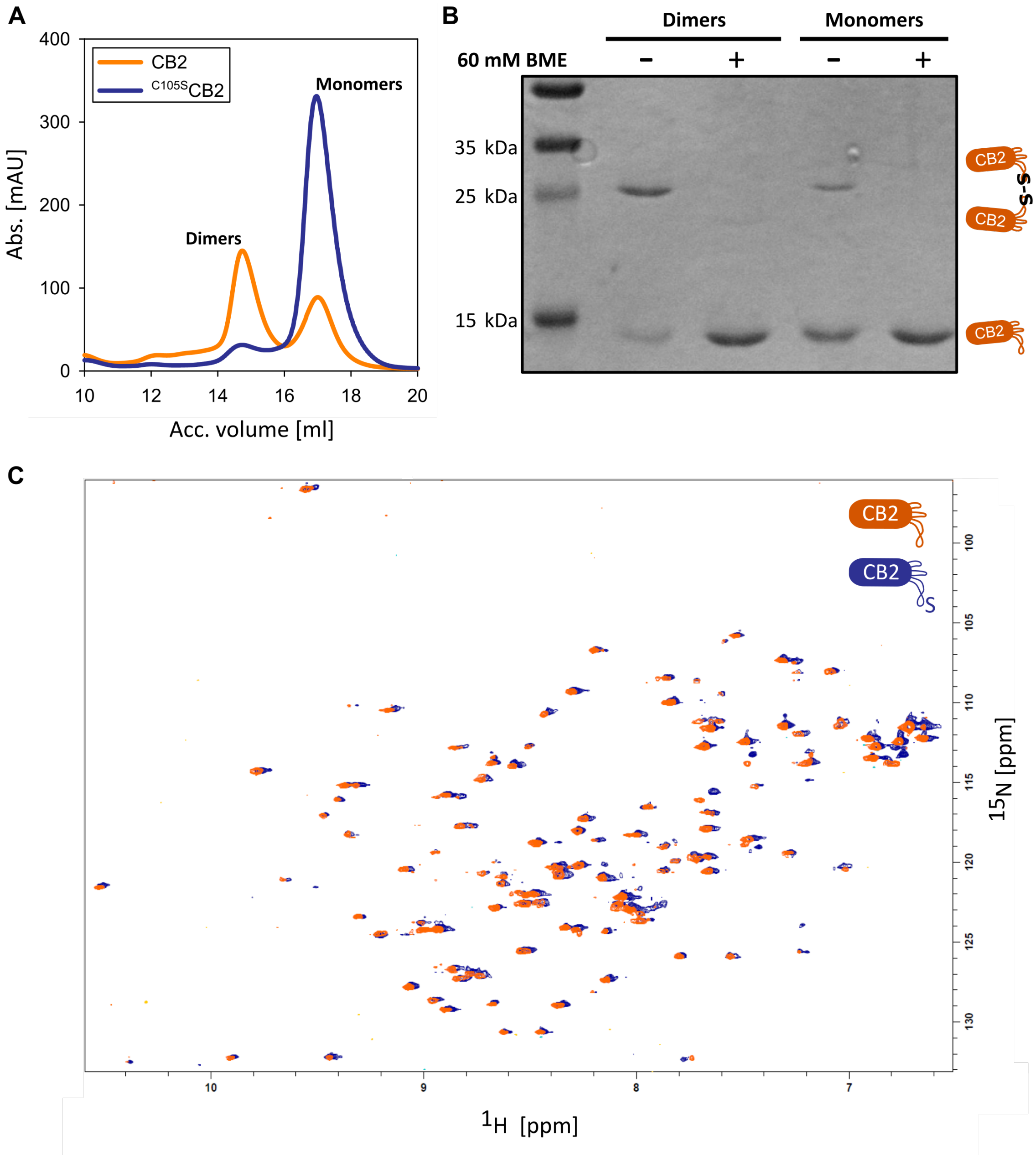


**Figure S2: Characterization of CB2 and $^{C105S}$CB2.** (A) Comparative size exclusion chromatography (SEC) of CB2 and $^{C105S}$CB2. Both proteins were purified as described in the method section but without DTT present. Subsequently, fractions were run on a Superdex™ 200 Increase FPLC column (23 ml) to compare elution profiles. Notably, a fraction of CB2 elutes earlier than the expected volume for a nanobody monomer (≈ 17 ml), indicating the presence of multimeric complexes. This additional peak is abolished in case of $^{C105S}$CB2. (B) Non-reductive SDS-PAGE of CB2 SEC fractions indicated in (A). Samples were incubated with native PAGE sample buffer with or without

60 mM beta-mercaptoethanol for 1 hour at 37°C and run on a standard SDS-PAGE gel. The gel shows that the additional SEC peak of CB2 represents dimers that are completely converted into monomers upon exposure to beta-mercaptoethanol. Replacement of cysteine-105 by serine prevents the dimerization. (C) Overlaid $^{1}H$–$^{15}N$ HSQC spectra of CB2 (orange) and $^{C105S}$CB2 (blue). No major signal shifts are visible, indicating the non-disruptive nature of the C105S mutation. CB2 and $^{C105S}$CB2 were acquired at 160 μM and 70 μM, respectively, in PBS buffer supplied with 0.75 mM DTT, 8 mM $NaN_3$, 0.5 mM DSS and 5% $D_2O$ at 298 K.

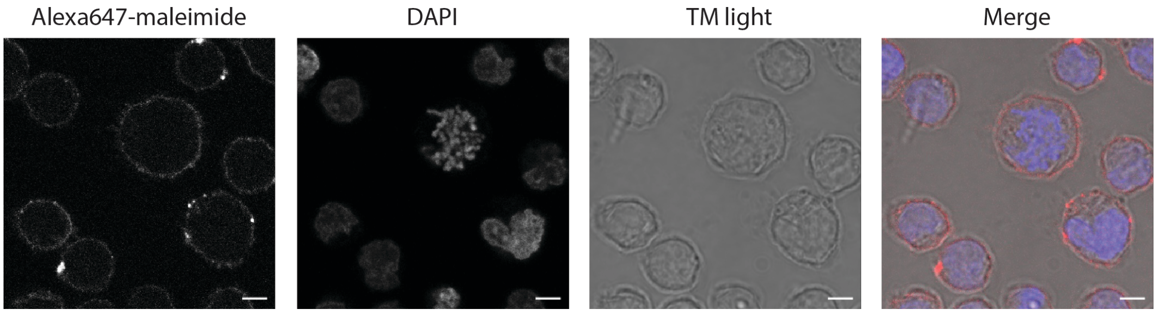


**Figure S3: Alexa647-maleimide stains cell surface thiols.** Confocal microscopy confirms that Alexa647-maleimide signal is only detected on the cell surface but not inside the cell. Colors in merge: red = Alexa647-maleimide, blue = DAPI. Scale bar = 5 μm.

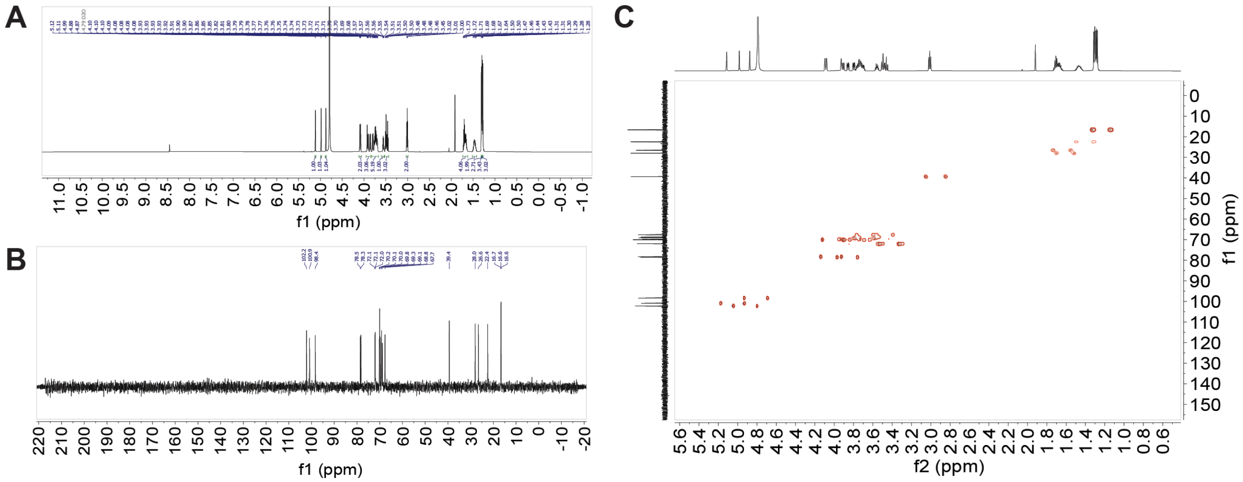


**Figure S4: NMR spectra of $Rha_3$.** (A) $^{1}H$ NMR (700 MHz, $D_2O$). (B) $^{13}C$ NMR (176 MHz, $D_2O$). (C) Coupled $^{13}C$,$^{1}H$ HSQC.

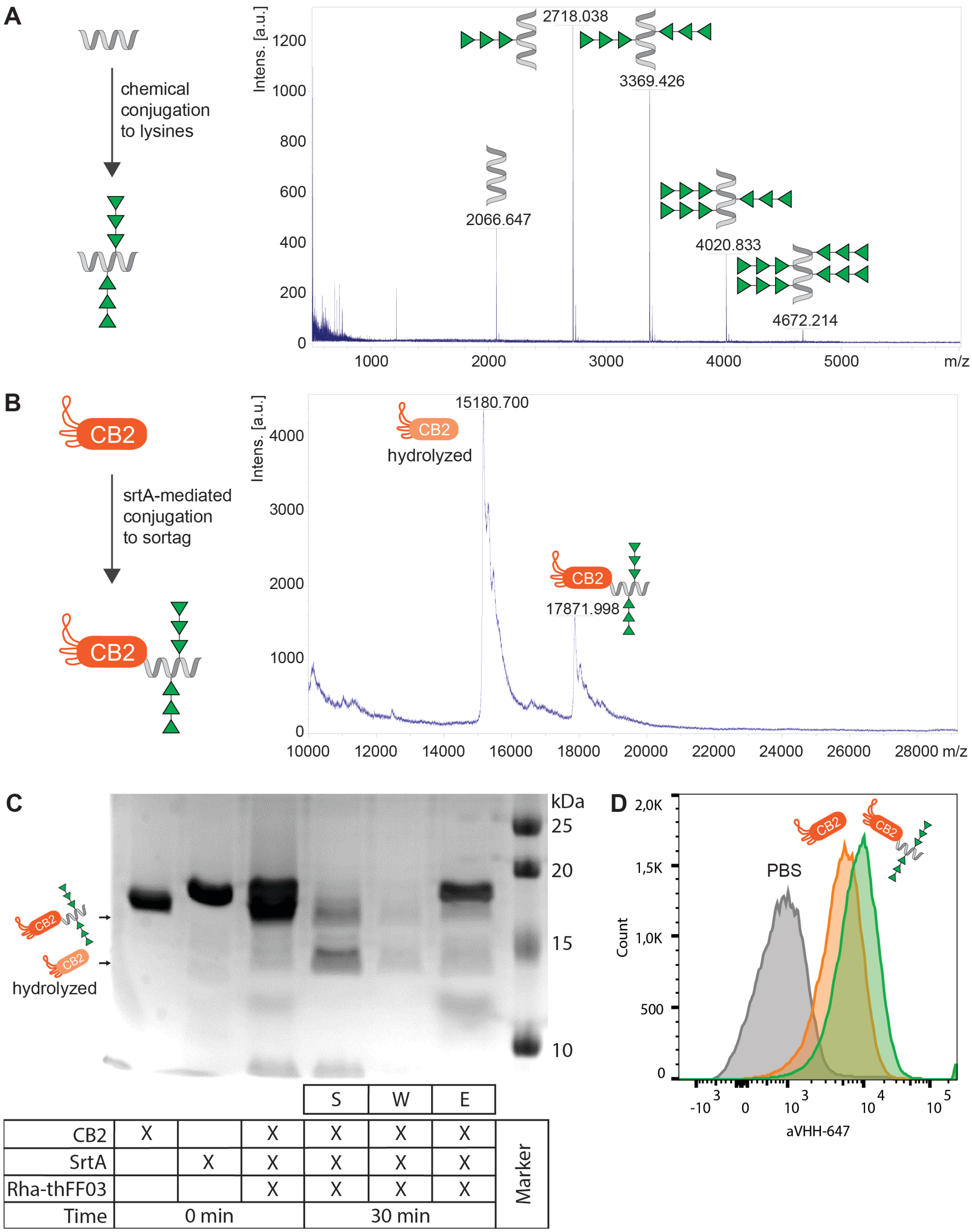
A
chemical conjugation to lysines
Intens. [a.u.]
2718.038
3369.426
2066.647
4020.833
4672.214
m/z
B
CB2
srtA-mediated conjugation to sortag
15180.700
hydrolyzed
17871.998
C
kDa
25
20
15
10
hydrolyzed
S
W
E
CB2
SrtA
Rha-thFF03
Time
0 min
30 min
Marker
D
PBS
Count
aVHH-647

**Figure S5: Conjugation of Rha-thFF03 to CB2.** (A) Chemical conjugation of $Rha_3$ to thFF03. The MALDI spectrum shows that 0-4 molecules of $Rha_3$ were deposited per peptide. (B-C) SrtA-mediated conjugation of Rha-thFF03 to CB2. (B) MALDI spectrum confirming the presence of CB2-Rha-thFF03 after the srtA reaction. (C) Coomassie-stained SDS-PAGE following the course of the srtA reaction over time. S, W and E correspond to supernatant, wash and eluate fractions when after 30 min of srtA reaction the conjugate is purified via Ni-NTA affinity chromatography. Note that both the desired product CB2-Rha-thFF03 (as expected) but also the hydrolyzed CB2 side product are found in the supernatant. (D) Flow cytometry histogram validating that CB2-Rha-thFF03 retains binding activity to SC-1 cells.

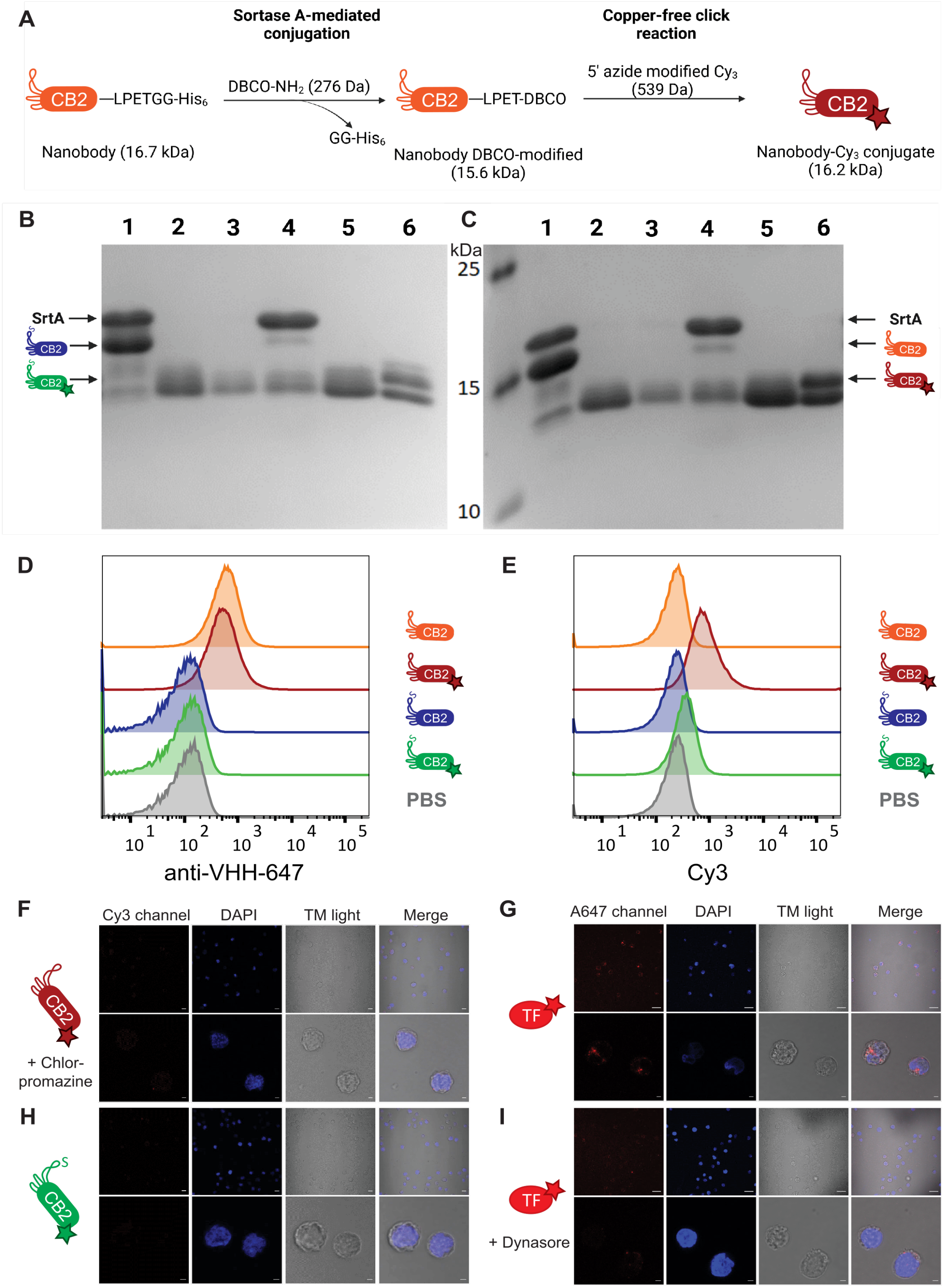

A
Sortase A-mediated conjugation
Copper-free click reaction
CB2 —LPETGG-His6
DBCO-NH2 (276 Da)
GG-His6
CB2 —LPET-DBCO
5' azide modified Cy3 (539 Da)
CB2
Nanobody (16.7 kDa)
Nanobody DBCO-modified (15.6 kDa)
Nanobody-Cy3 conjugate (16.2 kDa)
B
C
1 2 3 4 5 6
kDa
25
15
10
SrtA
CB2
D
E
anti-VHH-647
Cy3
PBS
F
Cy3 channel
DAPI
TM light
Merge
+ Chlor-promazine
G
A647 channel
TF
H
I
+ Dynasore

**Figure S6: Conjugation of Cy3 to CB2 and internalization controls.** (A) Schematic representation of the conjugation of Cy3 and the nanobody. At first, Nb was DBCO-modified via srtA-mediated conjugation and then a copper-free click reaction was used to conjugate Cy3. (B-C) Coomassie-stained SDS-PAGE gels of samples taken during the conjugation of Cy3 to (B) $^{C105S}$CB2 and (C) CB2. 1 = reaction mixture after 0 min; 2 = Ni-NTA flow-through containing DBCO-modified Nb; 3 = wash of Ni-NTA beads with srtA buffer; 4 = elution of remaining His-tagged Nb and srtA; 5 = pooled and concentrated DBCO-modified Nb after PD MidiTrap™ G-10 column; 6 = Nanobody Cy3 conjugate. (D-E) Binding to SC-1 cells in flow cytometry with unconjugated and Cy3-conjugated $^{C105S}$CB2, CB2 and PBS (negative control). No binding was detected for $^{C105S}$CB2 and $^{C105S}$CB2-Cy3. (F-I) Additional controls for CB2 internalization into SC-1 cells. Scale bar (first row) = 10 µm, scale bar (second row) = 2 µm. (F) CB2-Cy3 uptake in the presence of endocytosis inhibitor chlorpromazine. (G) Transferrin (TF) was used as a positive control for clathrin-mediated endocytosis. TF-Alexa647 uptake is visible in the red channel. (H) $^{C105S}$CB2-Cy3 was used as a negative control. Note that no signal is detected in the Cy3 channel with this protein. (I) TF-Alexa647 uptake in the presence of endocytosis inhibitor dynasore.

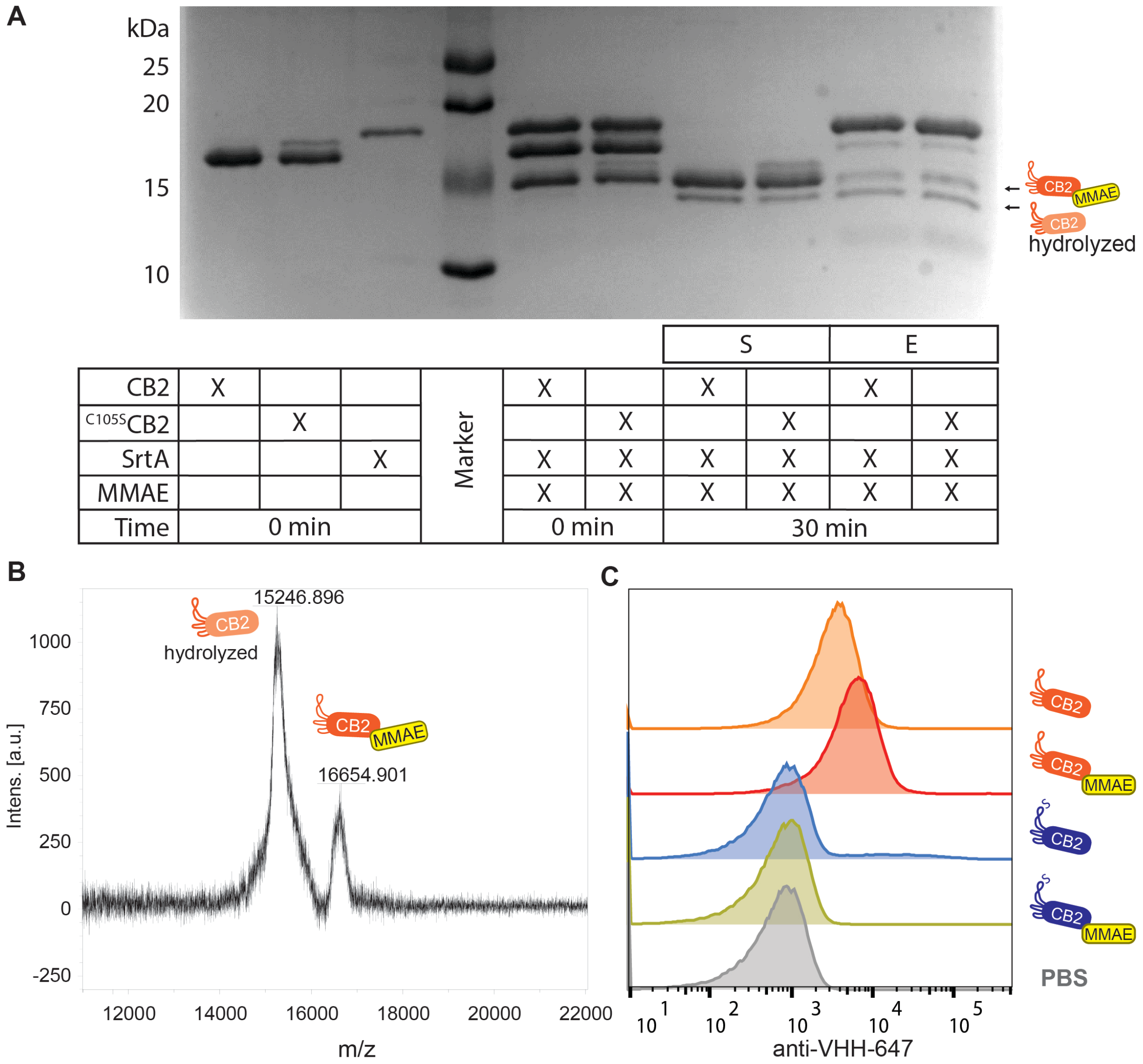


**Figure S7: Conjugation of MMAE to CB2.** (A) Coomassie-stained SDS-PAGE following the course of the srtA reaction over time. S and E correspond to supernatant and eluate fractions when after 30 min of srtA reaction the conjugate is purified via Ni-NTA affinity chromatography. The band intensity in the supernatant was quantified to determine the ratio of CB2-MMAE to hydrolyzed side product. (B) MALDI spectrum of the supernatant fraction from (A) confirming the presence of CB2-MMAE but also the hydrolyzed side product. (C) Flow cytometry histograms validating that CB2-MMAE retains full binding activity during conjugation. As expected, $^{C105S}$CB2-MMAE does not bind to SC-1 cells.
